# Intravenously Delivered Lipid Nanoparticles Access Acute Spinal Cord Injury via Disrupted Vasculature

**DOI:** 10.64898/2026.06.04.730155

**Authors:** Jacobus Burger, Luke Bolstad, Mia LaRico, Owen Lefebvre, Roy Ram Klein, Asma Eslami, William Murphy, Amgad Hanna, Daniel Hellenbrand

## Abstract

Trauma to the spinal cord disrupts the blood-spinal cord barrier and triggers a secondary injury cascade characterized by inflammation and progressive neuronal and glial cell death. Therapeutic cytokines and growth factors have shown promise as a treatment in preclinical studies, though their clinical translation is limited by short protein half-lives and the need for invasive intraspinal administration. Lipid nanoparticle-mediated delivery of mRNA offers an alternative strategy that enables transient protein production. Here, we investigated whether intravenously administered mRNA-lipid nanoparticles could leverage the injury-induced disruption of the blood-spinal cord barrier to access the injured spinal cord for local transgene expression. After spinal cord injury in a rat, lipid nanoparticles loaded with reporter mRNA were administered intravenously, and transgene expression was quantified in the spinal cord and peripheral organs. Intravenous delivery within a 6-hours post-injury resulted in local transgene expression in the injured spinal cord, demonstrating that mRNA-lipid nanoparticles cross the disrupted blood-spinal cord barrier. Transgene expression was observed in astrocytes, oligodendrocytes, microglia, and neurons, detected within 3 hours and remained elevated for up to 5 days post-injury. These findings demonstrate that systemic mRNA-lipid nanoparticles delivery exploit transient blood-spinal cord barrier disruption to achieve local gene expression in the injured spinal cord.

## Introduction

Spinal cord injuries (SCI) most commonly result from a traumatic fracture and/or dislocation of the spine, causing a compression, puncture or severing of the spinal cord within the spinal canal,^1,2^ which leads to paralysis below the level of injury and permanently compromising patient quality of life.^3–5^ Although significant advances have been made in instrumentation techniques for spinal column stabilization following fracture or dislocation, no clinically available treatments effectively reduce neuronal cell death or promote regeneration after SCI.^6,7^

The initial SCI trauma causes a disruption of the blood-spinal cord barrier (BSCB), initiating a secondary injury cascade that substantially expands the initial lesion.^8–10^ This cascade includes the recruitment and proliferation of resident microglia, extensive cytokine signaling, and infiltration of circulating immune cells.^8–11^ Together, oxidative stress, excitotoxicity, and inflammatory signaling further contribute to neuronal and glial cell death.^12,13^ As a result, functional neural tissue can be lost within minutes to hours following injury. Therefore, there has been an emphasis on developing therapeutic strategies capable of intervening during the acute phase of the injury response.

In animal models, several therapeutic approaches have been explored to mitigate the secondary injury cascade. Growth factors have been investigated to promote neuronal survival and regeneration in interneuron populations,^14–18^ while anti-inflammatory cytokines have been evaluated for their ability to attenuate the inflammatory environment that develops following injury.^19–25^ Although these strategies demonstrate strong therapeutic potential, significant translational challenges limit their clinical implementation.

Direct systemic administration of recombinant proteins can produce rapid biological effects; however, the short half-life of many cytokines and growth factors necessitates either repeated doses or administration of high systemic concentrations that can lead to unwanted off-target effects.^26–31^ Consequently, many preclinical studies rely on direct intraspinal injections to achieve localized delivery to the injury site.^20,23,25,32^ While promising results have been observed in animal studies, intraspinal injections require invasive surgical access to the spinal cord and are associated with risks including additional tissue damage, infection, and technical variability.^33–35^

In the clinical setting, preparation and administration of an intraspinal injection also require substantial time. Most patients will be initially transported to the nearest hospital, then transferred to a tertiary care center. Patients must first be hemodynamically stabilized and undergo imaging before surgery can be performed. As a result, injections would most likely occur during spinal stabilization surgery (>6 hours post-injury).^36,37^ This delay in administration misses the early therapeutic window, as inflammatory cytokine upregulation and neutrophil infiltration begin within the first 1–3 hours following SCI.^9,38,39^ These factors limit the translational feasibility of direct spinal cord injection for widespread clinical use.

Non-viral nucleic acid delivery systems offer an alternative approach that can overcome several spinal cord delivery limitations. Lipid nanoparticle (LNP) formulations have emerged as an effective platform for delivery of modified messenger RNA (mRNA-LNPs). mRNA-LNPs enable rapid cytoplasmic protein expression without requiring nuclear entry and exhibit transient expression profiles that may be advantageous when delivering potent cytokines or growth factors.^40–42^ In addition, the absence of viral components reduces the risk of vector-associated immunogenicity,^43–46^ while the use of modified mRNA vs wild type mRNA further reduces unwanted immune responses to foreign RNA.^47,48^

Importantly, traumatic SCI is accompanied by disruption of the BSCB, which increases vascular permeability at the injury site during the acute phase following trauma.^49–51^ This temporary barrier disruption presents a potential opportunity for systemically administered therapeutics to access the injured spinal cord. We hypothesized that injury-induced disruption of the BSCB would permit intravenously administered mRNA-LNPs to deliver mRNA to glial cells within the spinal cord during the acute phase of injury. Using a rat weight-drop contusion model of SCI, we characterized the systemic administration of mRNA-LNPs and evaluated reporter gene expression in the spinal cord and peripheral organs following injury. Results demonstrated that systemically delivered mRNA-LNPs effectively transfected central nervous system (CNS) cells within the injured spinal cord, with sustained transgene expression in the spinal cord for up to 5 days.

## Results

### Intravenously administered mRNA-LNPs crossed the disrupted BSCB and drove rapid transgene expression in the injured spinal cord

To evaluate whether intravenously administered mRNA-LNPs can cross the disrupted BSCB, LNP formulations were synthesized via microfluidics to generate nanoparticles approximately 140 nm in z-average diameter (**Fig. 1A-B**). Sprague Dawley rats implanted with jugular vein catheters underwent a T10 contusion injury followed by intravenous administration of LNPs loaded with firefly luciferase mRNA (Fluc mRNA-LNPs; **Fig. 1C**). To assess BSCB penetration, Fluc mRNA-LNPs were administered intravenously 0.5 hours after spinal cord contusion at a dose of 0.3 mg/kg mRNA. At 24 hours post-injury, rats were perfused with PBS to remove circulating blood, then spinal cords were harvested and incubated in 10 mg/mL luciferin prior to imaging using an in vivo imaging system (IVIS).

**Figure 1.**
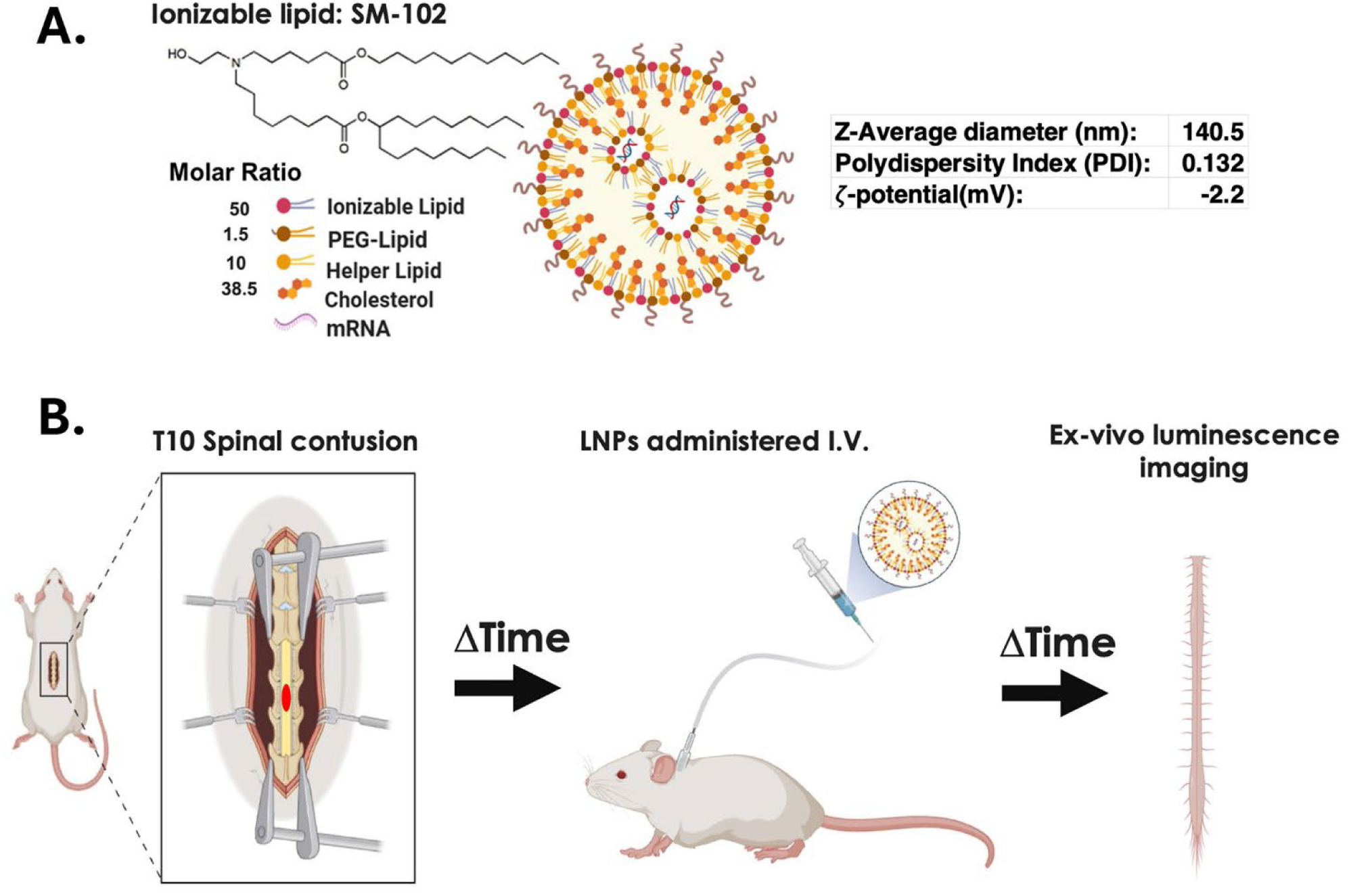
Experimental design for testing LNPs. LNPs were formulated using the indicated lipid component molar ratios, resulting in particles with an average diameter of 140.5 nm (**A**). To evaluate LNP efficacy, rats underwent a T10 spinal cord contusion, followed by administration of LNPs encapsulating firefly luciferase mRNA at 0.3 mg/kg mRNA via a jugular vein catheter (**B**). After the designated time interval, spinal cords were harvested, and luminescence was quantified *ex vivo* (**B**).

After SCI, Fluc mRNA-LNPs crossed the disrupted BSCB generating transgene expression at the injury site. When LNPs were administered intravenously after SCI, strong luminescent signal was detected at the injury site, indicating local luciferase expression in the injured spinal cord (**Fig. 2A**). In contrast, mRNA-LNPs did not transfect the spinal cord with intact BSCB. Intravenous administration of Fluc mRNA-LNPs in uninjured rats resulted in detectable luminescence in peripheral organs, predominantly the spleen, but no detectable signal was observed in the non-injured spinal cord (**Fig. 2B**) or brain (**Sup. Fig. 1**). These results demonstrated that mRNA-LNP entry into the CNS occurred through injury-induced disruption of the BSCB, and within the CNS transgene production remained spatially restricted to the lesion site.

**Figure 2.**
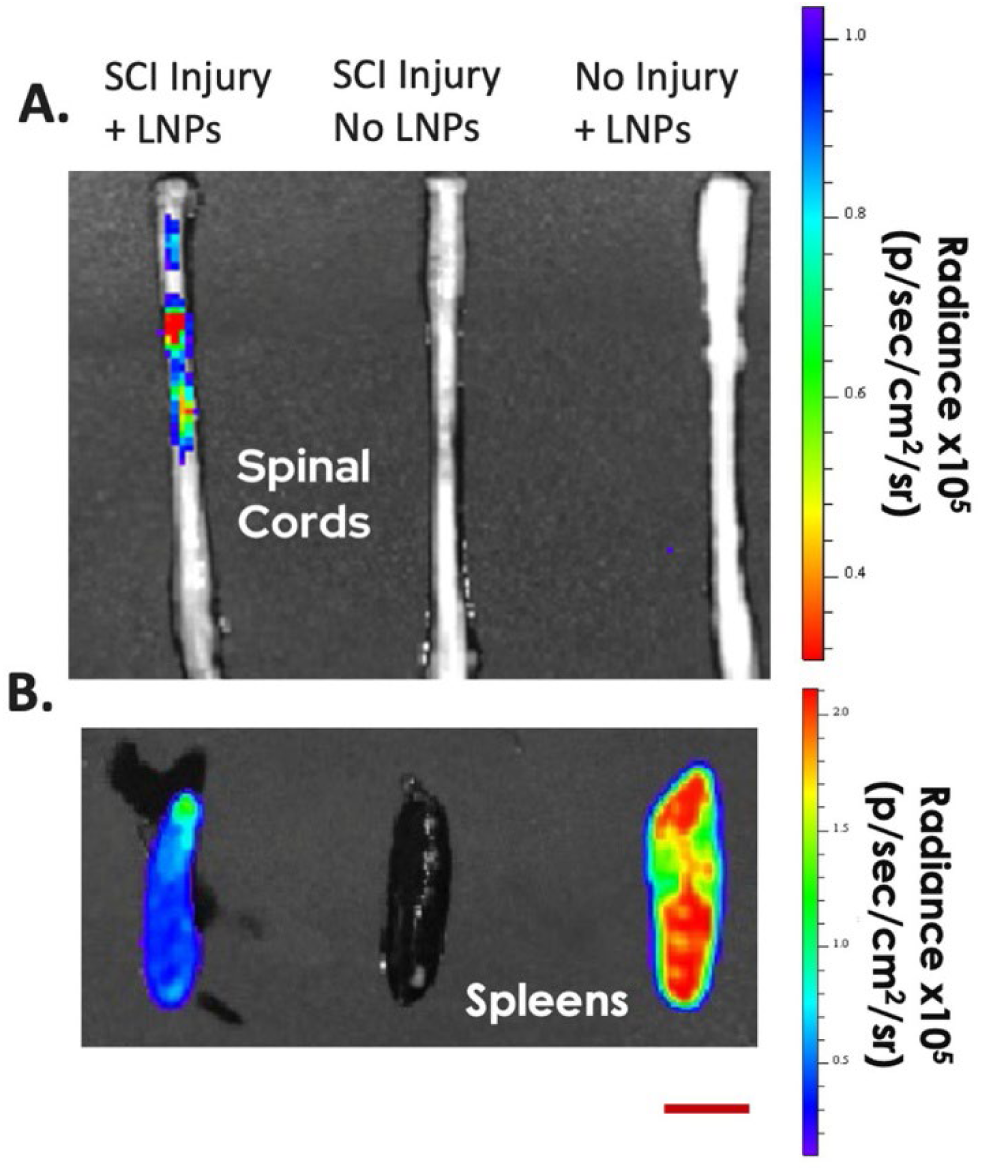
LNPs transfect the injured spinal cord. LNPs encapsulating Fluc mRNA were administered Intravenously 30 minutes after SCI, and luminescence was measured 24 hours post-injury. Following SCI, LNPs crossed the compromised BSCB and transfected cells within the injured spinal cord (**A**). In contrast, in uninjured animals, LNPs did not cross the intact BSCB (**A**). Intravenously administration of Fluc mRNA-loaded LNPs transfected the spleens in both SCI and uninjured rats (**B**). Scalebar = 10 mm.

The injury-induced disruption of the BSCB enabled a defined temporal window during which intravenously administered mRNA-LNPs could transfect cells in the injured spinal cord. To characterize this window, Fluc mRNA-LNPs were administered intravenously (0.3 mg/kg mRNA) through the jugular vein catheter at time points ranging from 0.5 hours to 12 hours following SCI. Luminescence imaging of spinal cords revealed the highest transfection when LNPs were administered 0.5 hours post-injury (**Fig. 3A-B**).

**Figure 3.**
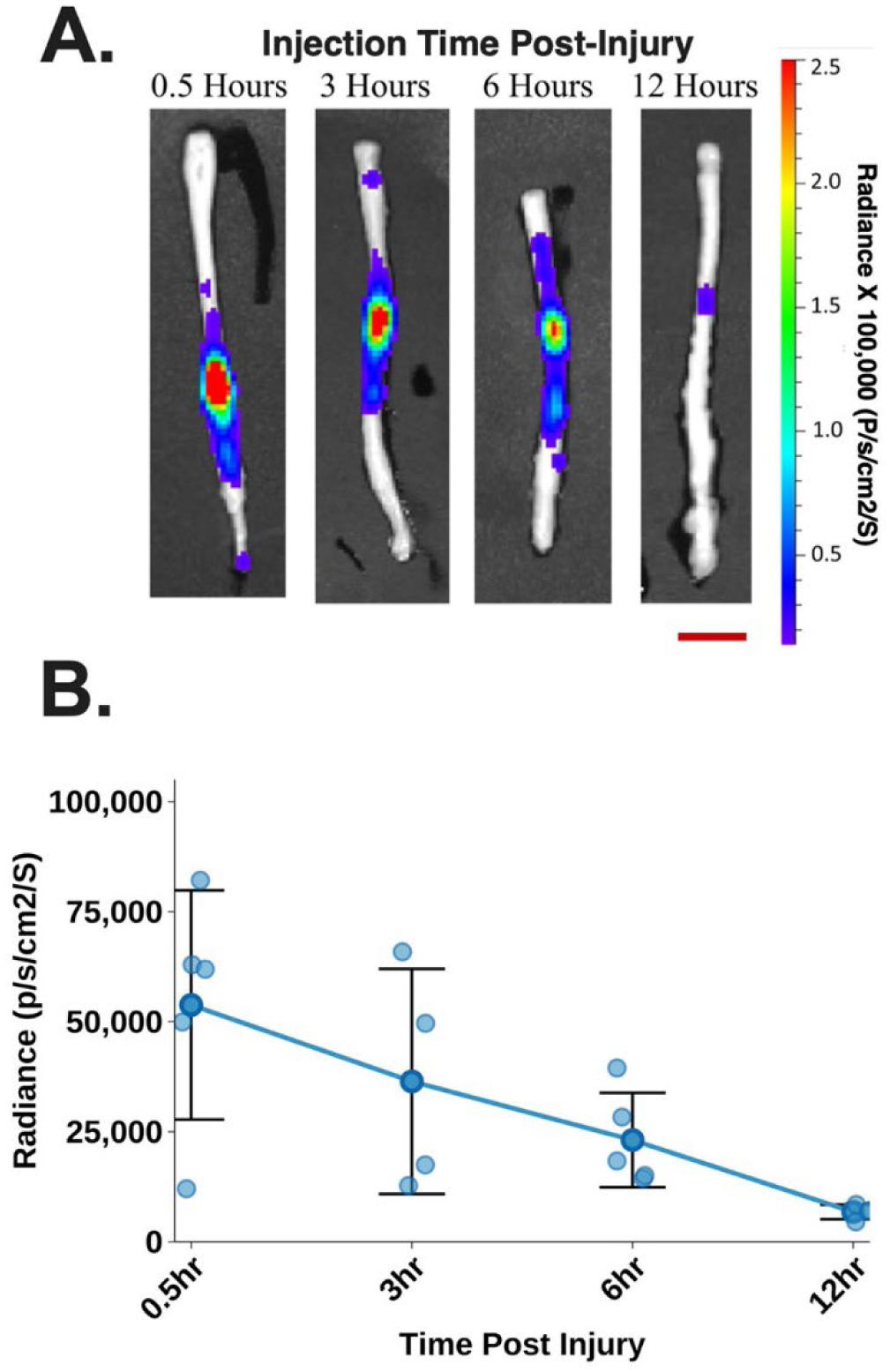
Timeline of BSCB disruption. Fluc mRNA-loaded LNPs were administered intravenously at varying times post-SCI and luminescence was measured 24 hours after LNP injection. IVIS images show efficient transfection in the spinal cord up to 6 hours post-injury (**A**). There was restoration of the BSCB over time with minimal transfection observed 12 hours post-SCI (**B**). Data are presented as mean ± SD; Scalebar = 10 mm. N = 5 for 0.5 hour and 6 hour groups. N = 4 for 3 hour and 12 hour groups.

Substantial transfection was still observed when LNPs were administered 3 hours post-SCI and 6 hours post-SCI. In contrast, administration at 12 hours post-injury resulted in minimal transfection, indicating substantial restoration of BSCB integrity by this time point (**Fig. 3A-B**). These data demonstrated that intravenous mRNA-LNP delivery enabled transfection within the injured spinal cord for at least 6 hours following SCI.

To determine which CNS cell types were transfected following delivery, Fluc mRNA-LNPs were administered intravenously 0.5 hours after spinal cord contusion (0.3 mg/kg mRNA). Spinal cords were harvested 48 hours post-SCI and analyzed by immunohistochemistry (IHC). Transverse spinal cord sections revealed a severe injury with central cord compression, in which the interior gray matter and the medial (inner) fibers of the white matter tracts were disrupted by the external force of the weight-drop contusion (**Sup. Fig. 2**).^2^ Luciferase-positive cells co-localizing with multiple CNS cell markers surrounding the lesion site, including O4⁺ oligodendrocytes and oligodendrocyte progenitor cells, Iba1⁺ microglia/macrophages, ACSA⁺ astrocytes, and NeuN⁺ neurons (**Fig. 4A-B**). More than 80% of the O4⁺ cells and ACSA⁺ cells were transfected and expressed luciferase (**Fig. 4B**). These findings indicate that intravenously delivered mRNA-LNPs transfected multiple CNS cell types within the injured spinal cord, with high transfection of astrocyte and oligodendrocyte lineage cells.

**Figure 4.**
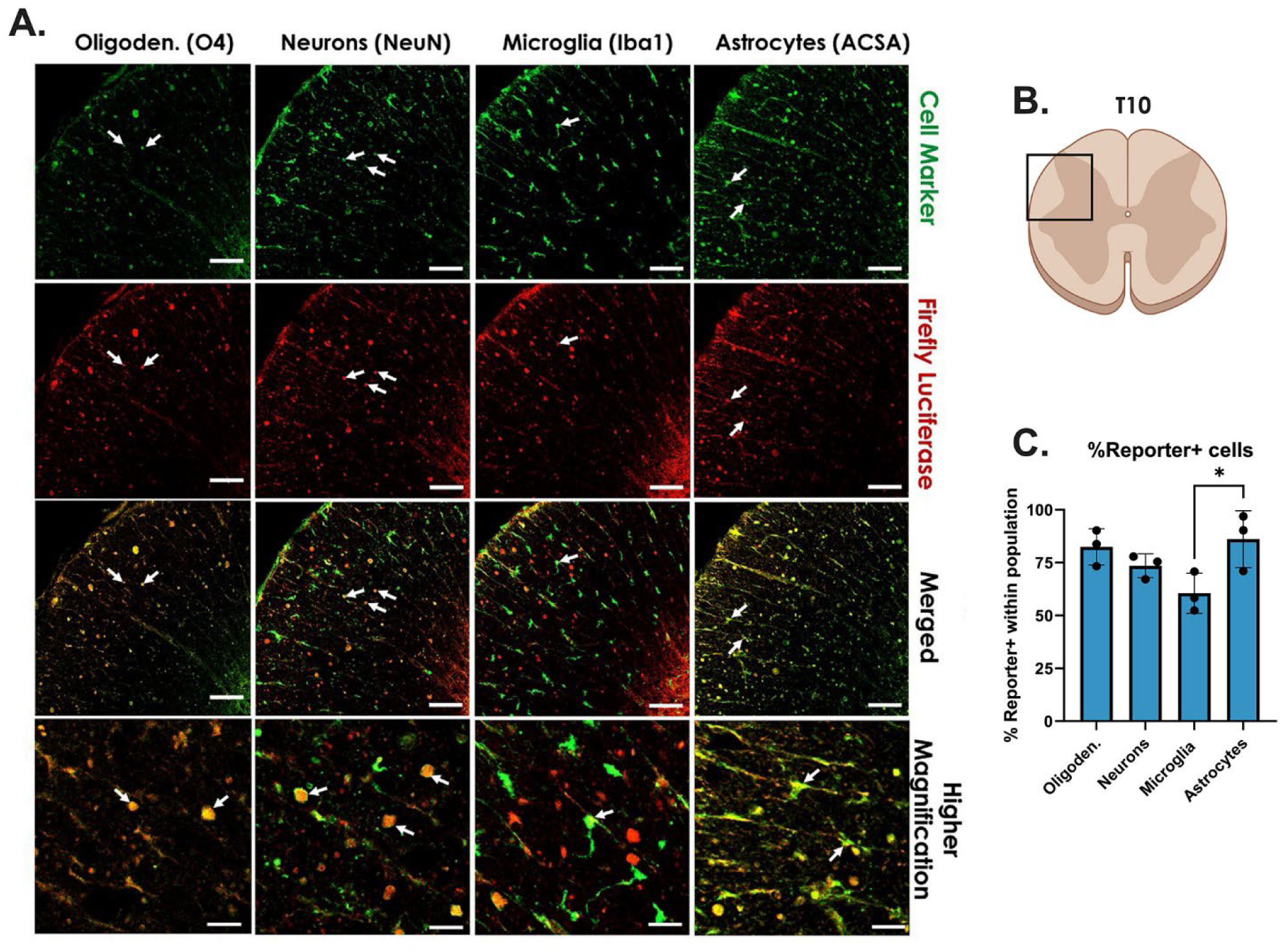
Intravenously administered LNPs transfect CNS cells within the injured spinal cord. IHC was performed to label oligodendrocytes, neurons, microglia, and astrocytes (**A**) in transverse spinal cord sections that were harvested 48 hours post-SCI (**B**). The number of marker⁺ cells within each cell type population co-expressing reporter signal was quantified within the same field of view, and the percent positive within each cell type population was calculated across three biological replicate tissue sections within the ventral horn(**C**). Arrows point to cell types expressing the Luciferase transgene. Data are presented as mean ± SEM; *P <0.05 by one-way ANOVA with Tukey’s post hoc analysis. Scalebars = 100 μm, for Higher magnification images scalebars = 40 μm.

To characterize the temporal profile of mRNA translation, Fluc mRNA-LNPs were administered intravenously 0.5 hours after SCI. Spinal cords were harvested between 3 hours and 120 hours post-SCI and luminescence was measured with IVIS. Luciferase expression was elevated at the earliest time assessed, 3 hours post-injection (**Fig. 5A-B**). Luciferase expression peaked at 24 hours post-SCI and remained elevated through 72 hours post-injury before declining with minimal transgene expression by 120 hours post-SCI (**Fig. 5A-B**). These results demonstrated rapid uptake of mRNA-LNPs into the injured spinal cord followed by transient transgene expression at the lesion site.

**Figure 5.**
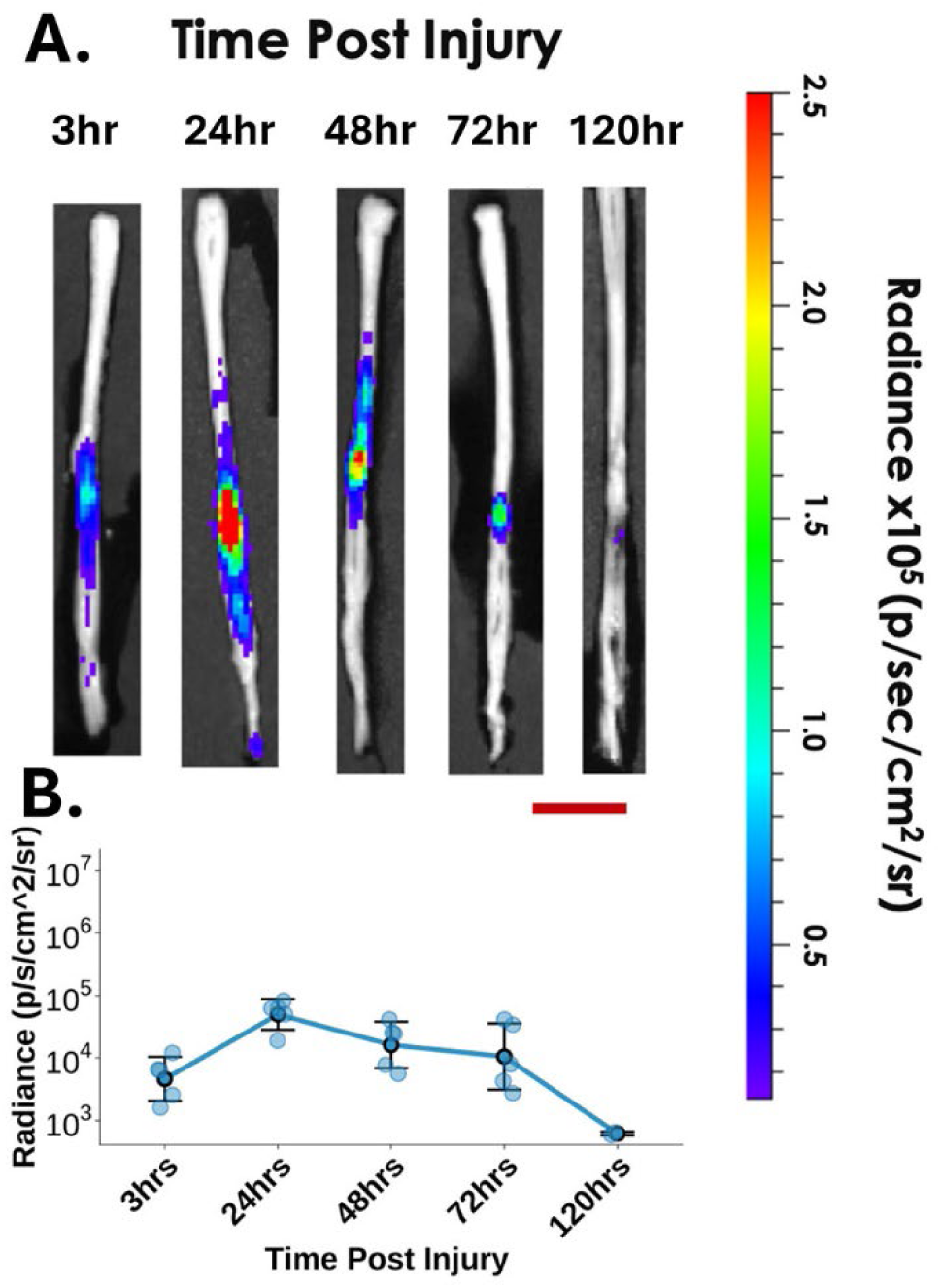
mRNA translation profile. Fluc mRNA-loaded LNPs were administered Intravenously through a JVC 0.5 hours post-SCI, and luminescence was measured at varying times after injury. IVIS images show efficient luciferase mRNA translation in the spinal cord starting as early as 3 hours post-injury and continuing for 120 hours (**A**). The luciferase mRNA translation peaked 24 hours post-injury and there was minimal translation 120 hours post-injury (**B**). Data are presented as mean ± SD. N = 4 for 120 hours post-injury, N = 5 all other times post-injury. Scalebar = 10 mm.

### Intravenously administered mRNA-LNPs drove transgene expression in the liver, lungs, and spleen

Previous studies have shown that intravenously administered LNPs transfect the lungs, liver, and spleen.^52–54^ Therefore, we quantified luciferase expression in these organs following intravenous administration of Fluc mRNA-LNPs 0.5 hours after SCI. In the lungs, luciferase expression was elevated 3 hours post-SCI, peaked at 24 hours post-SCI, and then declined rapidly, with minimal expression detected by 48 hours post-SCI (**Fig. 6A**). In the liver luciferase expression was already near peak levels by 3 hours post-injury and remained at a high level for 24 hours, before declining substantially by 48 hours (**Fig. 6B**). Similarly, in the spleen luciferase expression was near peak levels by 3 hours post-injury and remained at a high level for 24 hours (**Fig. 6C**). Luciferase expression in the spleen subsequently declined steadily until there was minimal transgene expression by 120 hours post-SCI (**Fig. 6C**). Taken together, these results indicated rapid uptake of mRNA-LNPs into the lungs, liver, and spleen followed by transient transgene expression in each organ.

**Figure 6.**
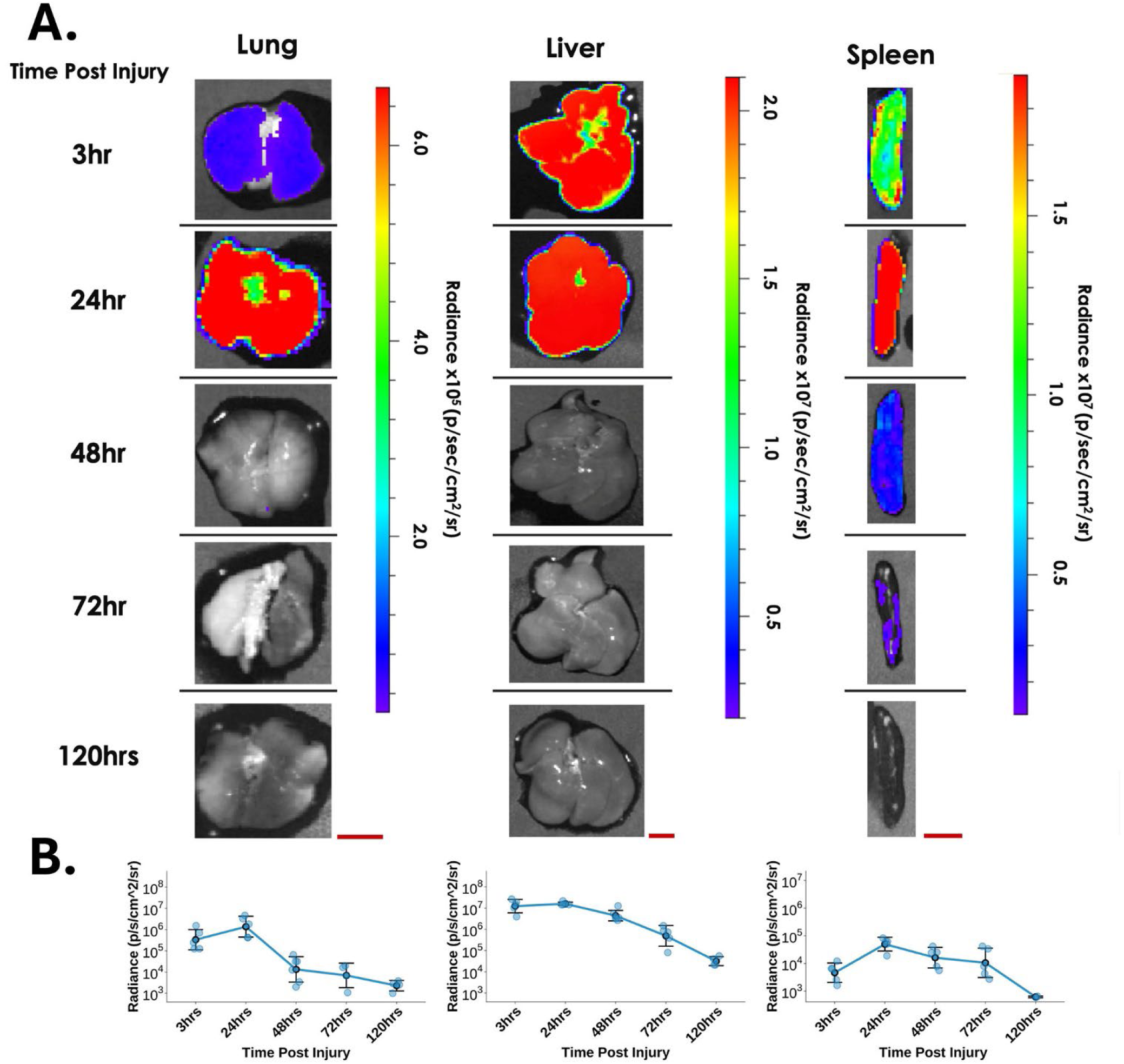
Translation of Fluc mRNA in peripheral organs. Fluc mRNA-loaded LNPs transfected the lungs, liver, and spleen following intravenous administration of mRNA-LNPs 0.5 hours post injury (**A**). Transgene expression was quantified for 120 hours post injury (**B**). Data are presented as mean ± SD. N = 4 for 120 hours post-injury, N = 5 all other times post-injury. Scalebars = 10 mm.

## Discussion

SCI causes cell death and disruption of the BSCB, initiating a secondary injury cascade leading to a cytokine storm and extensive infiltration of immune cells.^8–10^ Although there are positive effects of inflammation after SCI, the extensive infiltration of immune cells contributes to additional neural degeneration.^12,13^ Therefore much research has worked to develop drug delivery platforms to deliver treatments to reduce inflammation after SCI.

In this study, the mechanical disruption of the BSCB following SCI created a temporally restricted window that enabled intravenously administered mRNA-LNPs to access the lesion site and promoted transgene expression in the injured spinal cord. Imaging of the reporter gene delivered revealed maximal spinal cord transfection when LNPs were administered within the first 6 hours post-injury, followed by a rapid decline in signal as barrier integrity was restored. These findings align with previously reported timelines of BSCB disruption following contusion injury and demonstrated that this transient vascular permeability can be exploited for nucleic acid delivery to the injured spinal cord.^55,56^ To our knowledge, this work provides the first demonstration that systemically administered mRNA-LNPs can leverage injury-induced BSCB disruption to achieve local gene delivery within the CNS during the acute phase of SCI.

Lipid nanoparticles enabled efficient systemic delivery of mRNA to the injured spinal cord without the need for invasive intraparenchymal injections. Several experimental gene therapy strategies for SCI rely on viral vectors delivered directly into spinal cord tissue,^32,57–61^ which introduces surgical complexity and risks additional tissue damage. Viral systems also exhibit delayed expression kinetics due to the requirement for nuclear entry and transcription prior to protein production.^43,62^ In contrast, mRNA-LNPs mediate rapid cytoplasmic translation and transient expression, allowing therapeutic proteins to be produced within hours following administration.^43^ The rapid transgene expression observed in the injured spinal cord in this study may be particularly advantageous for therapies targeting acute inflammatory responses after SCI, as the first 12 hours following injury are characterized by substantial production of pro-inflammatory cytokines and infiltration of immune cells.^9,11,38,39^ Therefore, systemic delivery of mRNA-LNPs represents a potential alternative to repeated intraspinal injections of recombinant proteins or viral-based CNS gene therapy approaches.

The mRNA-LNP approach produced functional proteins directly by cells within the injured spinal cord microenvironment. Cell-type analysis indicated that transgene expression occurred across multiple CNS populations, including oligodendrocyte-lineage and astrocyte cells surrounding the lesion site.

Importantly, the O4 antibody labels both oligodendrocytes and, to an even greater extent, oligodendrocyte progenitor cells,^63,64^ which proliferate extensively and accumulate in large numbers within the astrocytic border after SCI.^65,66^ Local production of functional proteins by resident cells may be particularly useful for modulating the post-injury inflammatory response. For example, anti-inflammatory cytokines expressed directly within the lesion microenvironment may directly influence macrophage polarization to mitigate inflammation and promote tissue repair. ^21,67,68^ Delivering mRNA to cells within the lesion site may therefore enable functional proteins to act precisely where they are most biologically relevant.

In principle, the strategy described here may also extend beyond SCI to other neurological pathologies associated with vascular disruption. Traumatic brain injury (TBI) and hemorrhagic stroke are characterized by acute breakdown of the blood-brain barrier within affected tissue, similar to SCI.^69–71^ Recently, in a controlled cortical impact model of TBI in mice, LNPs were shown to transfect cells in the injured brain for up to 48 hours post-injury.^72^ The longer duration of access to damaged brain tissue may be due to differences between TBI and SCI, or it may reflect greater injury severity in the TBI model compared to the SCI model used. It is also possible that the smaller LNPs (∼100 nm) used in the TBI model were able to traverse the disrupted vasculature over a longer timeframe.^72^ However, a previous study evaluating the effects of nanoparticle size following TBI, observed a significant reduction in accumulation of 100 nm polystyrene beads within injured brain tissue between 1 and 6 hours post-injury.^56^ These finding suggests that nanoparticle composition and surface charge may also play critical roles in uptake.^73^ Regardless, transient permeability of the neurovascular barrier permits systemically delivered mRNA-LNPs to access damaged regions of the brain. Therefore, exploiting injury-induced barrier disruption represents a broader strategy for achieving local gene delivery across multiple acute CNS insults.

While there was efficient transfection of multiple cell types in the spinal cord, systemic mRNA-LNP administration also resulted in substantial transgene expression within the lung, liver, and spleen. These findings are consistent with the well-characterized biodistribution of LNP formulations following intravenous administration.^52,74^ Similar to previous studies, the high levels of luciferase produced in the liver occurred over a short duration with most of the translation ceasing 48 hours after systemic LNP administration.^52^ Although peripheral expression declined within days, off-target transgene production in these organs may be undesirable for certain therapeutic payloads. Future studies should therefore focus on strategies to reduce peripheral expression while preserving delivery to the injury site. LNPs are highly adaptable, and modifications to surface targeting moieties can be made to increase transfection in specific organs or cells of interest.^53,75–77^ Another potential approach is the incorporation of tissue-specific RNA regulatory elements within the therapeutic mRNA to selectively suppress translation in off-target organs such as the liver.^78^ Such regulatory elements have been previously shown to effectively restrict expression of systemically delivered mRNA therapeutics while maintaining expression in target tissues.^79^ Taken together, non-viral LNP-based mRNA delivery offers several conceptual advantages over protein and other nucleic acid-based approaches with the ability to target a specific tissue type.

In this study, mRNA-LNPs loaded with Fluc mRNA (0.3 mg/kg mRNA; ∼3 mg/kg lipid) sustained transgene expression in the injured spinal cord for 5 days.^80^ In certain treatment paradigms for SCI, it may be necessary to maintain transgene expression for a longer timeframe. If a more sustained or amplified gene delivery is desired, a self amplifying or circular RNA modality could also further improve on-target protein production.^81,82^ These constructs allow for augmented gene replacement therapy via increased construct stability (Circular) and construct replication (self-amplifying). Combining these regulatory strategies with injury-localized delivery may further improve the safety profile of mRNA-LNP-based therapeutics for neurotrauma applications.

Although this work demonstrated the ability of LNPs to cross the disrupted BSCB and transfect cells within the injured spinal cord, this study was limited by the use of a single LNP formulation employing SM-102 as the ionizable lipid. The choice of ionizable lipid is a key determinant of LNP organ and cell-type tropism following intravenous administration, and alternative ionizable lipids may yield different biodistribution profiles or altered targeting efficiency to the injured spinal cord.^53,83^ Future studies should evaluate additional LNP formulations with distinct ionizable lipids to determine whether the injury-induced BSCB disruption can be leveraged across a broader range of delivery systems, and to identify formulations that further enhance on-target delivery to the SCI lesion while minimizing peripheral organ expression.

## Conclusions

In summary, our findings demonstrate that intravenously administered mRNA-LNPs can exploit injury-induced disruption of the BSCB to achieve local gene expression within the acutely injured spinal cord. This strategy enabled rapid and transient delivery of functional proteins during the early phase of injury without requiring invasive procedures. Because this approach relies on intravenous administration within a defined acute therapeutic window, it may be compatible with early intervention following traumatic injury, including administration prior to arrival at a tertiary care center. Leveraging injury-induced vascular permeability for intravenous mRNA delivery may therefore represent a promising strategy for the development of gene-based therapeutics targeting acute CNS injuries.

## Materials and Methods

### Lipid nanoparticle formulation and physicochemical characterization

An ethanol phase containing an ionizable lipid (SM-102, Medkoo Biosciences, Durham, NC, USA), structural lipid, cholesterol, and pegylated lipid were prepared at lipid component molar ratios of 50, 10, 38.5 and 1.5 respectively (**Fig. 1A**). An aqueous phase containing the payload mRNA was prepared by diluting 1 mg/ml mRNA in 10mM citrate buffer. Ethanol and aqueous phases were mixed using a microfluidic device. The ratio of lipids to nucleic acid payload was determined by maintaining a set ratio of ionizable nitrogen in the lipids to anionic phosphates in the mRNA backbone of 6:1.

SM-102 was selected as the ionizable lipid because it is a clinically validated component of the Moderna COVID-19 mRNA vaccine (mRNA-1273) and has demonstrated efficient mRNA delivery following intravenous administration across numerous preclinical and clinical studies. The use of an approved formulation component also aligns with emerging FDA guidance encouraging sponsors to leverage prior manufacturing knowledge and established safety profiles of related products to streamline regulatory development of future mRNA-based therapeutics.

Lipid nanoparticle hydrodynamic diameter and size distribution were measured using dynamic light scattering (DLS). LNP formulations were diluted in RNase-free water or phosphate-buffered saline (PBS) to an appropriate scattering concentration. Measurements were performed at 25 °C using a fixed-angle light scattering detector. For each sample, multiple sequential acquisitions were collected and averaged to ensure reproducibility.

Particle size was reported as the z-average hydrodynamic diameter (nm). The polydispersity index (PDI) was calculated from the same cumulant fit and used as a measure of particle size distribution uniformity. For each formulation, at least three independent measurements were performed and the mean values of z-average size and PDI were reported. LNP preparations were accepted for experimental use only where PDI < 0.2 and z-average diameter <200nm (**Fig. 1B**).

### Animal model & surgical procedure

All surgical procedures were performed in accordance with the National Institutes of Health Guide for the Care and Use of Laboratory Animals and protocols approved by the institution’s Animal Care and Use Committee. Analyses were performed *ex vivo* on the spinal cord, lungs, liver, and spleen. To ensure consistency in organ size, all animals used were female Sprague Dawley rats (10-12 weeks old; approximately 225 g; Envigo, Huntingdon, UK). All rats receiving intravenous treatments were purchased with a pre-implanted jugular vein catheter.

For spinal cord contusion surgeries, anesthesia was induced using ketamine (90 mg/kg) and xylazine (7 mg/kg) administered via intraperitoneal injection (Midwest Veterinary Supply Inc., Lakeville, MN, USA) and a laminectomy was performed at the T10-T11 vertebral level to expose the dura covering the spinal cord. The rat was stabilized and slightly elevated above the platform using clamps placed on the T9 and T12 spinous processes. A MASCIS Weight Drop Device (WM Keck Center for Collaborative Neuroscience, Rutgers University, Piscataway, NJ, USA; Model II) was used to induce spinal cord contusion by dropping a 10 g weight from a height of 25 mm ^25,84^. Postoperative analgesia consisted of slow-release buprenorphine (1.2 mg/kg) administered subcutaneously (ZooPharm, LLC, Laramie, WY, USA).

At predetermined times post-injury, LNPs loaded with firefly luciferase mRNA (TriLink BioTechnologies, San Diego, USA) were administered via the jugular vein catheter. The lowest of three previously tested doses was used (0.3 mg/kg mRNA; ∼3 mg/kg lipid) ^80^.

Following surgery, rats were housed two per cage at room temperature under a standard light-dark cycle and were provided an antibiotic-enriched diet (Uniprim; Envigo) for one week. Bladders were manually expressed at least twice daily until tissue collection to prevent urinary tract infection. Animals that developed urinary tract infections were treated with enrofloxacin (10 mg/kg) administered subcutaneously once daily for four consecutive days. Rats were also closely monitored for dehydration and received subcutaneous saline when signs of dehydration were observed.

For tissue collection, rats were euthanized with a lethal dose of isoflurane and transcardially perfused with cold PBS to remove blood prior to harvesting the spinal cord and other organs. For luminescence measurements, spinal cords and organs were incubated in 10 mg/mL luciferin (Gold Biotechnology, Inc., St. Louis, MO, USA) for 15 minutes before imaging with an in vivo imaging system (IVIS; **Fig. 1C**). (60 second exposure).

### Immunohistochemistry (IHC) cell-type transgene expression analysis

Harvested spinal cords were fixed in 4% paraformaldehyde (PFA) for 24 hours, followed by cryoprotection in 20% sucrose for 24 hours and then in 30% sucrose for at least 24 hours. A 3 mm spinal cord segment centered on the injury site was embedded in Tissue-Tek OCT compound, frozen, and sectioned transversely at a thickness of 20 µm. Sections were rinsed in 0.1 M phosphate-buffered saline (PBS) for 10 minutes and then incubated in blocking solution (4% donkey serum, 1% bovine serum albumin (BSA), and 0.5% Triton X-100 in 0.1 M PBS) for 1 hour at room temperature. After rinsing, sections were incubated overnight at 4°C with primary antibodies diluted in 0.1 M PBS containing 1% BSA. The following day, sections were washed three times in 0.1 M PBS (5 minutes each) and incubated with secondary antibodies for 2 hours at room temperature in the dark.

Primary antibodies were used at the following dilutions: goat anti-luciferase IgG (1:250, Abcam, Cambridge, United Kingdom), rabbit anti-Iba1 (1:500, Abcam), mouse anti-O4 (1:250, R&D Systems, Minneapolis, MN, USA), mouse anti-ACSA-2 biotinylated (1:250, Miltenyi Biotec, Bergisch Gladbach, Germany), and mouse anti-NeuN (0.1 µg/mL, Abcam).

Secondary antibodies (Thermo Fisher Scientific, Waltham, MA, USA) were used at the following dilutions: donkey anti-goat Alexa Fluor 594 (1:500), donkey anti-rabbit Alexa Fluor 488 (1:500), donkey anti-mouse Alexa Fluor 488 (1:500), and streptavidin 488 (1:500). Nuclei were counterstained with 4’,6-diamidino-2-phenylindole (DAPI).

Confocal microscopy was used to quantify reporter gene expression in oligodendrocytes, neurons, microglia, and astrocytes. Nuclei were first identified in the DAPI channel using automated threshold-based segmentation to detect individual cells. The centroid of each segmented nucleus was used as the reference position for cell-associated fluorescence measurements. Cell-type markers were identified using ACSA-2, NeuN, Iba1, and O4 staining, while reporter expression was detected in the luciferase channel. Cells were classified as marker-positive or reporter-positive based on fluorescence intensity above background levels. Reporter positivity was calculated as the proportion of marker-positive cells exhibiting detectable reporter signal within the same field of view. Quantification was performed on single-field images acquired at 10× magnification using automated image segmentation and intensity-based classification.

### Statistical Analysis

Prism 10.4.0 (GraphPad Software, San Diego, CA, USA) was used for data analysis, statistics, and generating graphs. Figure legends include number of animals per treatment group, statistical and post hoc tests, and degree of significance for each comparison. Differences were considered significant at P < 0.05. All quantitative data are presented as mean ± standard deviation. All schematics were created with BioRender.com.

## Data availability

The datasets used and analyzed during the current study are included within the article and its supplemental files. Any related data will be made available on request.

## Acknowledgements

This work was supported by the National Institutes of Health grant R01 NS136564-01. The authors would also like to acknowledge the University of Wisconsin Small Animal Imaging & Radiotherapy Facility (SAIRF), University of Wisconsin Translational Research Initiatives in Pathology (TRIP) Laboratory, and the UW Carbone Cancer Center Support Grant (NCI P30 CA014520) for providing pathology services, the IVIS Spectrum imaging, and technical support for this study.

## Author Contributions

Conceptualization: J.C.B., D.J.H.; methodology: J.C.B., L.J.B., D.J.H.; investigation: J.C.B., L.J.B., D.J.H.; data curation: J.C.B., L.J.B., M.J.L., O.R.L., R.R.K., A.E.,; visualization: J.C.B., L.J.B., D.J.H.; funding acquisition: W.L.M., A.S.H.; supervision: W.L.M., A.S.H., D.J.H.; writing – original draft: J.C.B., D.J.H.; writing – review and editing: J.C.B., L.J.B., W.L.M., A.S.H., D.J.H.;

## Declaration of interests

W.L.M. is a co-founder and Chief Science Officer at Stem Pharm, Inc. and Dianomi Inc. The other authors declare that they have no competing interests.

---

**Supplementary Figure 1.**
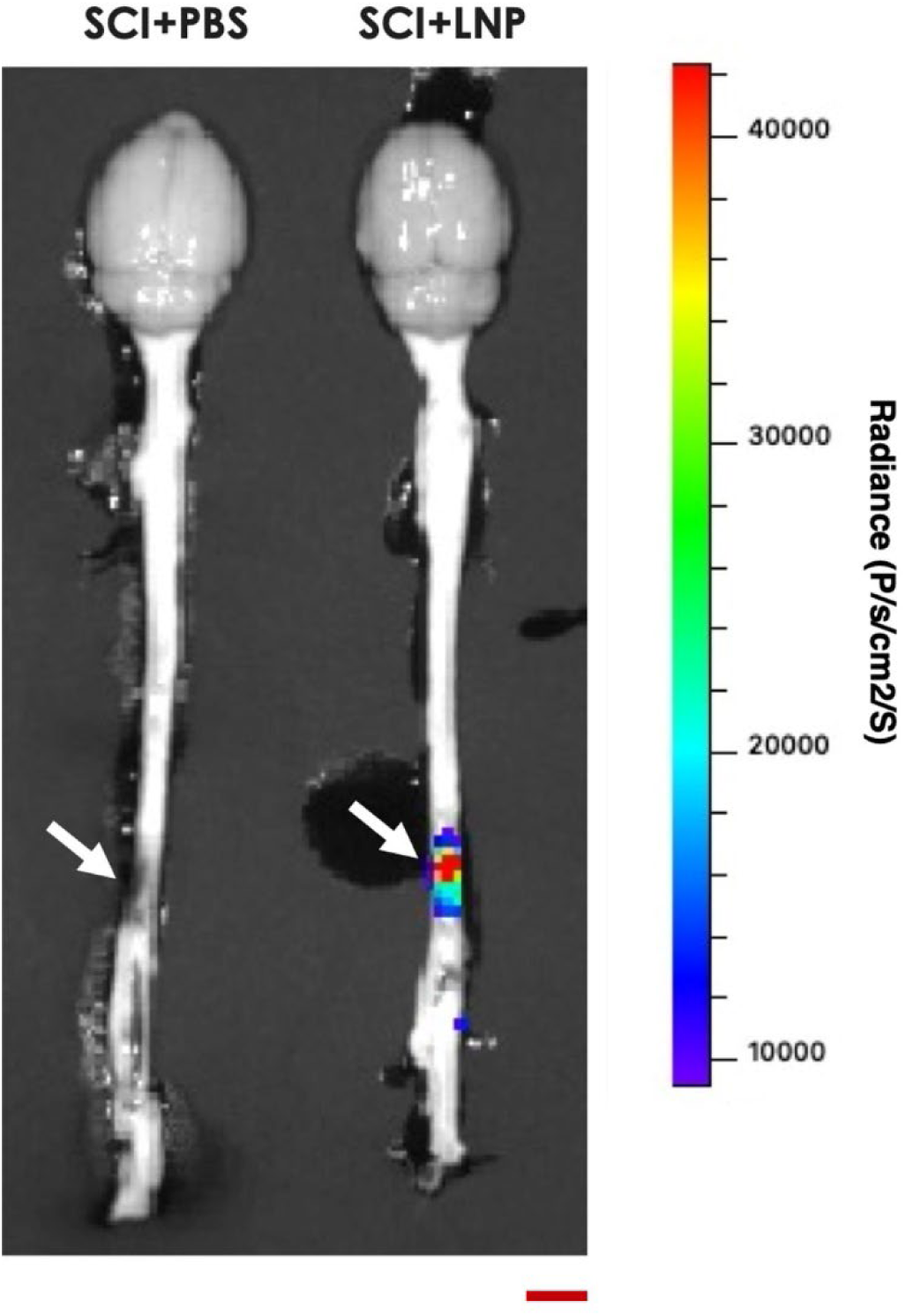
LNPs do not cross the intact BSCB or the blood-brain barrier. Rats were subjected to a T10 SCI, and Fluc mRNA-loaded LNPs were injected via a JVC 30 minutes post-injury and the entire brainstems were harvested 24 hours post-injury and imaged for bioluminescence. White arrows point to T10 contusion site. Scalebar = 10 mm.

**Supplementary Figure 2.**
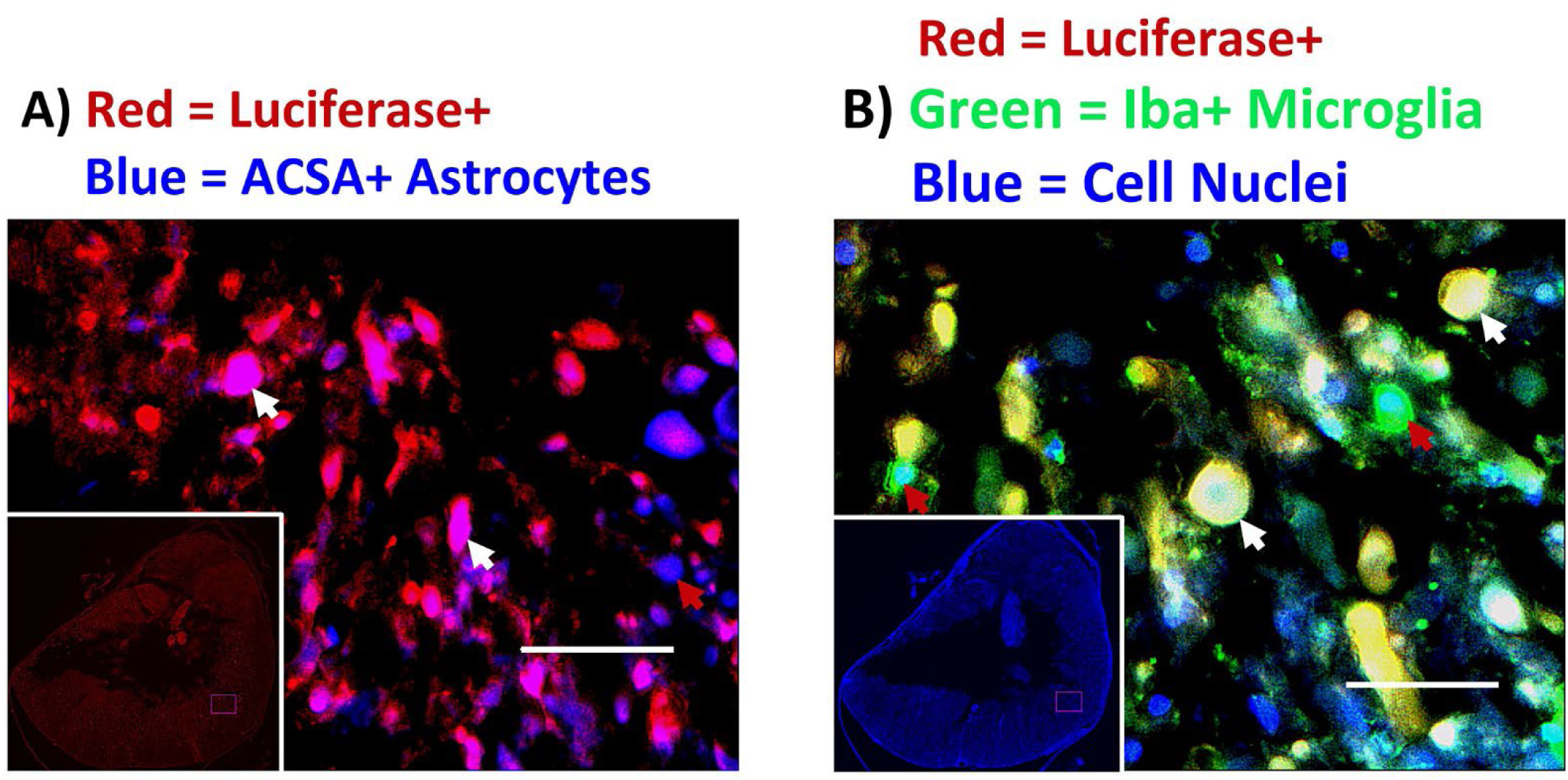
LNPs transfect cells surrounding the disrupted interior gray matter and the medial (inner) fibers of the white matter tracts. Rats were subjected to a T10 SCI, and Fluc mRNA-loaded LNPs were injected via a JVC 30 minutes post-injury. The spinal cords were then harvested 24 hours post-injury. Transfected astrocytes (A) and microglia (B) surrounded the entire periphery of the injury epicenter on transverse sections from T10 spinal cord. White arrows point to transfected cells and red arrows point to cells that are not transfected. Scalebars = 50 µm.

